# Content-based dissociation of hippocampal involvement in prediction

**DOI:** 10.1101/568303

**Authors:** Peter Kok, Lindsay I. Rait, Nicholas B. Turk-Browne

## Abstract

It has recently become clear that one of the key functions of the hippocampus is to predict future inputs. In line with this, previous research has revealed prediction-related signals in the hippocampus for complex visual objects, such as fractals and abstract shapes. Based on this, it has been suggested that the hippocampus may generate perceptual expectations, especially when relying on rapidly learned predictive associations between arbitrary stimuli. However, it is currently unknown whether the hippocampus implements general-purpose computations that subserve all associative predictions, regardless of stimulus properties, or whether the involvement of the hippocampus is stimulus-dependent. To investigate this, we exposed male and female human participants to complex auditory cues predicting either the shape of a complex object (Experiment 1) or the orientation of a simple line grating (Experiment 2). We measured brain activity using high-resolution functional magnetic resonance imaging (fMRI), in combination with inverted encoding models to reconstruct shape and orientation representations in visual cortex and the hippocampus. Our results revealed that expectations about shape and orientation evoked distinct representations in the hippocampus. For complex shapes, the hippocampus represented which shape was expected, potentially serving as a source of top-down predictions. In contrast, for simple gratings, the hippocampus represented only unexpected orientations, more reminiscent of a prediction *error.* We discuss several potential explanations for this dissociation, concluding that the computational role of the hippocampus in predictive processing depends upon the nature and complexity of stimuli.

**Significance Statement:** To deal with the noisy and ambiguous sensory signals received by our brain, it is crucial to use prior knowledge of the world to guide perception. Previous research suggests that the hippocampus is involved in predicting upcoming visual stimuli based on prior knowledge. In the current study, we show that hippocampal prediction is specific to expectations of complex objects, whereas for simple features the hippocampus generates an opposite prediction error signal instead. These findings demonstrate that the computational role of the hippocampus can be content-sensitive and refine our understanding of the involvement of memory systems in predictive processing.

## Introduction

Sensory processing is strongly influenced by prior expectations (1). Expectations about both simple features (e.g., orientation) (2–4) and more complex objects (e.g., shape) (5–9) modulate processing in visual cortex. However, it is unclear whether these two kinds of expectations arise from the same top-down sources and operate via the same underlying mechanisms.

Previous research has revealed prediction-related signals in the hippocampus for complex visual objects, such as fractals (10, 11) and abstract shapes (12, 13). These studies used fMRI to reveal that the pattern of activity in the hippocampus contains information about expected visual objects, upon presentation of a predictive cue. Based on this, it has been suggested that the hippocampus may generate perceptual expectations, especially when these predictions result from rapidly learned associations between arbitrary stimuli (11, 14–16). The role of the hippocampus may be particularly relevant when the associations are cross-modal, given its bidirectional connectivity with all sensory systems (17).

This perspective raises the possibility that the hippocampus implements general-purpose computations that subserve all kinds of (associative) prediction (18–20). That is, upon presentation of a predictive cue or context, the hippocampus may retrieve the associated outcome through pattern completion (14, 21), regardless of the exact nature of the stimuli. This is in line with evidence that the hippocampus is involved in many different types of predictions, pertaining to, for example, faces and scenes (22), auditory sequences (23), odors (24), and spatial locations (19, 25).

However, an alternative and untested hypothesis is that the hippocampus only generates certain types of predictions. Theories casting sensory processing as hierarchical Bayesian inference (26–28) suggest that each brain region provides predictions only to those lower-order region(s) with which it has direct feedback connections, rather than bridging the full hierarchy. For instance, V2 may supply V1 with predictions about the locations of short line elements making up longer lines (26, 29). Visual processing in areas of the medial temporal lobe (MTL) most directly connected to the hippocampus, such as perirhinal and parahippocampal cortices, is dominated by high-level objects and scenes, respectively (30–32), as well as their spatial, temporal, and associative relations (33–35). Processing in these regions is thought to be abstracted away from low-level sensory features (17, 31), such as orientation and pitch. Because of this high-level selectivity of MTL cortex, hippocampal predictions may only traffic in complex visual stimuli.

To distinguish these hypotheses, we conducted two nearly identical studies that varied only in the complexity of visual predictions. We exposed human participants to complex auditory cues predicting either the shape of an abstract Fourier descriptor (Experiment 1, N=24) or the orientation of a Gabor grating (Experiment 2, N=24). We measured brain activity using high-resolution fMRI, and used inverted encoding models (36) to reconstruct shape and orientation information from neural representations in visual cortex and the hippocampus. To preview, we found that expectations about both orientation and shape modulated visual cortex, facilitating processing of expected stimuli relative to unexpected stimuli. However, expectations about orientation and shape were represented very differently in the hippocampus.

## Results

Participants were exposed to auditory cues that predicted either which complex shape was likely to be presented (Experiment 1, Fig. 1a-d), or the likely orientation of an upcoming grating stimulus (Experiment 2, Fig. 1e-h). In both experiments, two stimuli were presented on each trial, which were either identical, or slightly different from one another (Experiment 1: second shape slightly warped, Experiment 2: second grating slightly different phase). Participants were asked to report whether the two stimuli on any given trial were the same or different.

**Figure 1.**
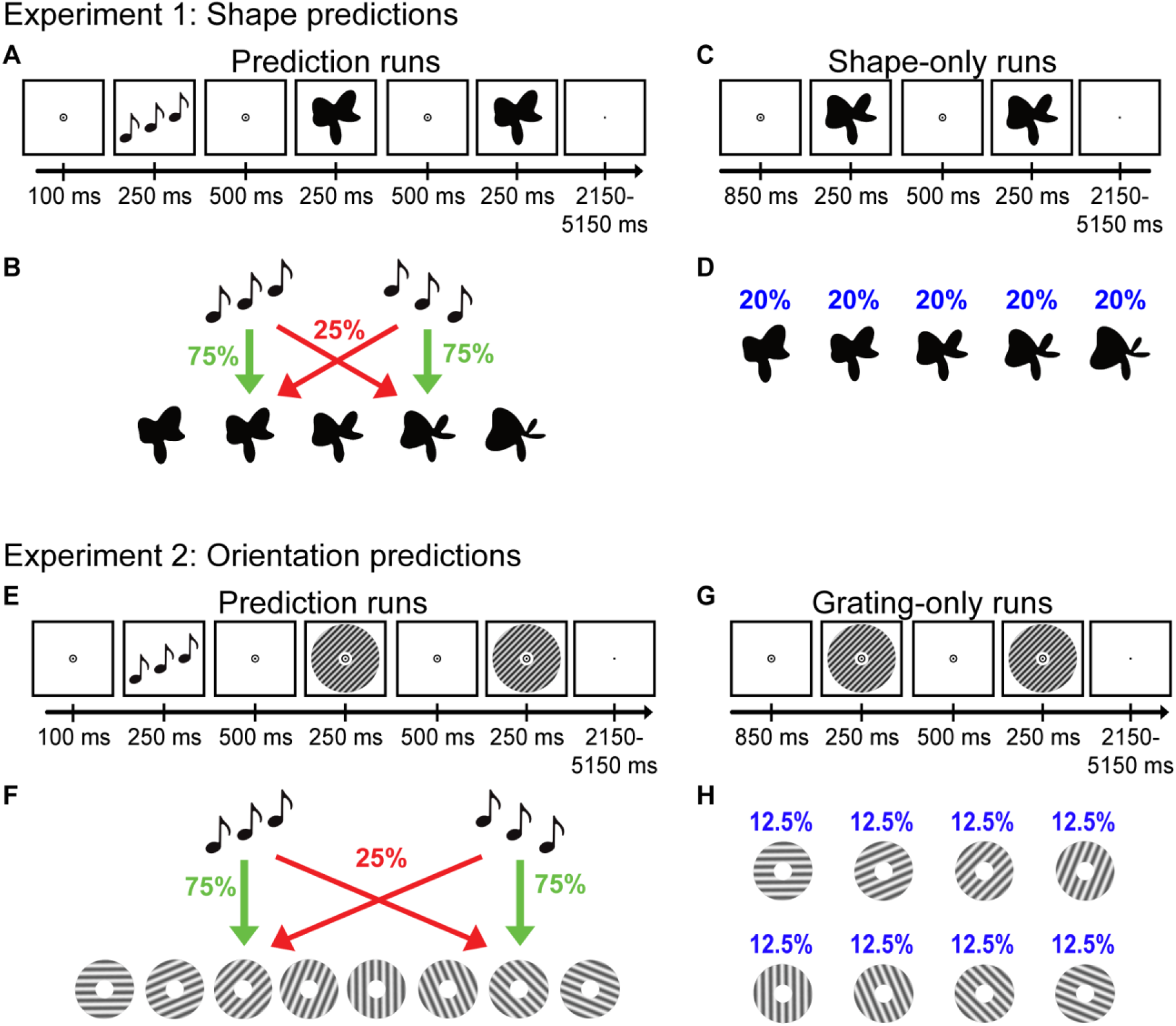
Experimental paradigms. ***A***, During prediction runs in Experiment 1, an auditory cue preceded the presentation of two consecutive shape stimuli, where the second shape was either identical to the first or slightly warped. Participants indicated whether the two shapes were the same or different. ***B***, The auditory cue (ascending vs. descending tones) predicted whether the first shape would be shape 2 or shape 4 (of 5 shapes). The cue was valid on 75% of trials, whereas in the other 25% of (invalid) trials the other, unpredicted shape was presented (prediction runs) or the shapes were omitted (omission runs). ***C***, During shape-only runs, no auditory cues were presented. As in the other runs, two shapes appeared sequentially and participants reported same or different. ***D***, All five shapes appeared with equal (20%) probability on trials of the shape-only runs. ***E***, The design of Experiment 2 was identical to Experiment 1, except that the shapes were replaced by oriented gratings. The second grating was either the same as the first or its phase was shifted slightly. ***F***, The auditory cue predicted whether the first grating would be rotated 45° or 135°, and this cue was valid 75% of the time. ***G***, During grating-only runs, no predictive auditory cues were presented. ***H***, All eight gratings were equally likely to appear (12.5%) during grating-only runs.

### Behavioural results

Participants were able to discriminate small differences in both complex shapes (36.9 +/− 2.3% modulation of the 3.18 Hz radial frequency component, mean +/− SEM) and simple gratings (2.7 +/− 0.2 radians phase difference, mean +/− SEM) during the visual stimuli only runs. This was also the case during the prediction runs for both complex shapes (valid trials, 31.6 +/− 2.5%; invalid trials, 33.2 +/− 2.9%) and simple gratings (valid trials, 2.4 +/− 0.2 radians; invalid trials, 2.4 +/− 0.2 radians). For Experiment 1, accuracy and reaction times (RTs) did not differ between valid trials (accuracy, 70.6 +/− 1.2%; RT, 575 +/− 16 ms) and invalid trials (accuracy, 68.8 +/− 1.5%; RT, 573 +/− 18 ms; both *p* values > 0.20). In Experiment 2, participants were slightly more accurate for valid (75.2 +/− 1.3%) than invalid (72.0 +/− 1.5%) trials (*t*_(23)_ = 2.25 *p* = 0.03). RTs did not differ significantly between conditions (valid: 646 +/− 13ms; invalid: 646 +/− 14ms, *p* = 0.93). Since the phase differences between the two gratings on valid and invalid trials, respectively, were controlled by separate staircases, we checked that this difference in accuracy was not accompanied by a significant difference in the staircase threshold (i.e., task difficulty) between the two conditions (*t*_(23)_ = 0.36, *p* = 0.72).

### Visual cortex

As expected, the pattern of activity in visual cortex contained a representation of the shape (Experiment 1, V1: *t*_(23)_ = 14.72, *p* = 3.4 x 10^-13^; V2: *t*_(23)_ = 14.23, *p* = 6.8 x 10^-13^; LO: *t*_(23)_ = 7.04, *p* = 3.5 x 10^-7^) or grating orientation (Experiment 2, V1: *t*_(23)_ = 11.32, *p* = 7.1 x 10^-11^; V2: *t*_(23)_ = 13.48, *p* = 2.1 x 10^-12^; LO: *t*_(23)_ = 8.28, *p* = 2.4 x 10^-8^) that was presented on the screen. Interestingly, the temporal evolution of these representations was strongly affected by the auditory prediction cues (Fig. 3). We characterized the time courses of the decoding signal by fitting a canonical (double-gamma) hemodynamic response function (HRF) and its temporal derivative. The parameter estimate of the canonical HRF indicates the peak amplitude of the signal, whereas the temporal derivative parameter estimate reflects the latency of the signal (37, 38).

**Figure 2.**
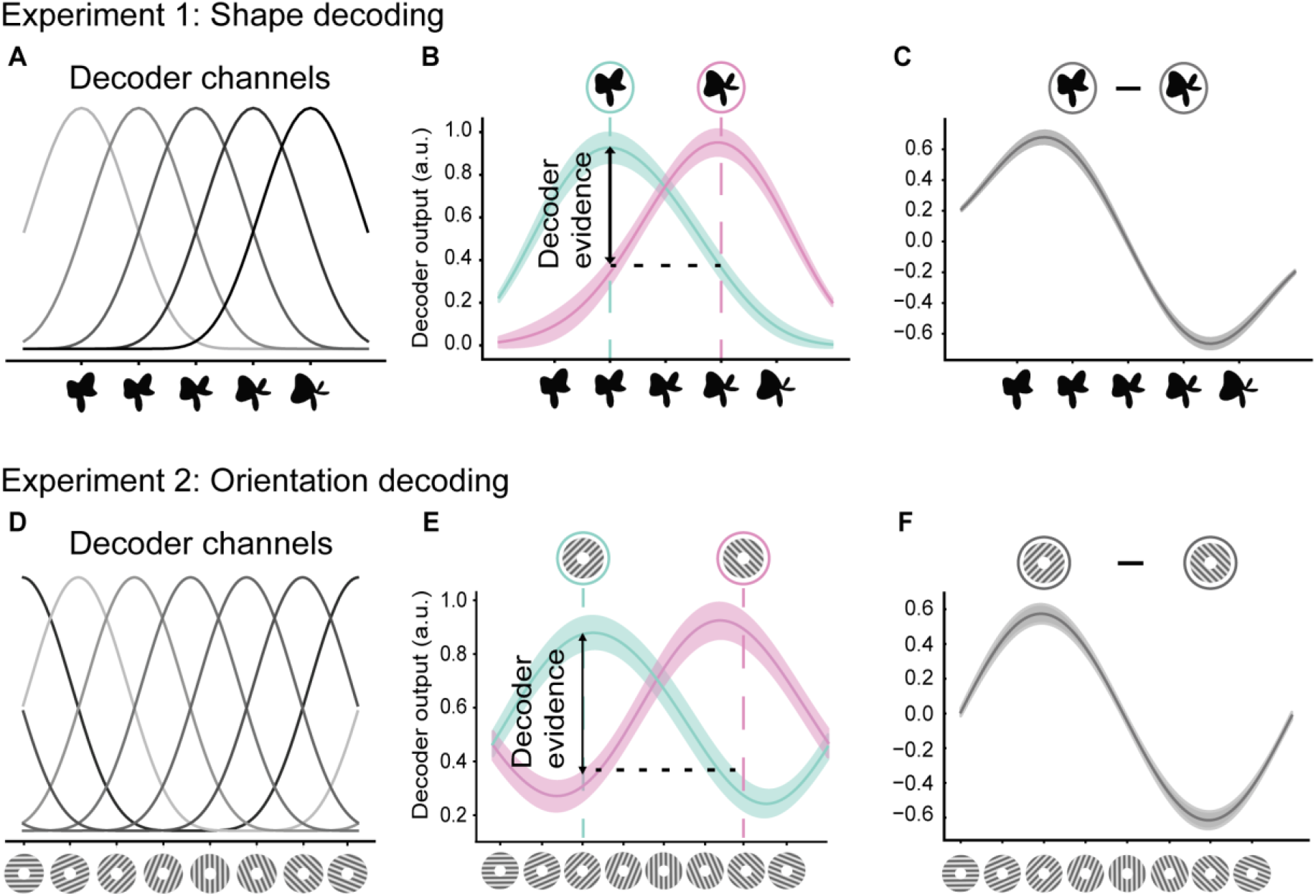
Illustration of the decoding methods. ***A***, We used a forward modelling approach to reconstruct shapes from the pattern of BOLD activity. Shape selectivity was characterized by five hypothetical channels, each with an idealized shape tuning curve. BOLD patterns obtained from the shape-only runs were used to estimate the weights on the five hypothetical channels separately for each voxel, using linear regression. ***B***, Using these weights, the second stage of the analysis reconstructed the channel outputs associated with the pattern of activity across voxels evoked by the prediction and omission runs (only shapes 2 and 4 were used in these runs). Channel outputs were converted to a weighted average of the five basis functions, resulting in neural evidence across shape space. Decoding performance was quantified by subtracting the evidence at the presented shape (e.g., shape 2) from the evidence at the non-presented shape (e.g., shape 4). ***C***, Finally, solely for the purpose of visualising the shape-specific information, we collapsed across the presented shapes by subtracting the neural evidence for shape 4 from that for shape 2, thereby removing any non-shape-specific BOLD signals. ***D***, An identical forward modelling approach was used in Experiment 2, except that orientation selectivity was characterized by six hypothetical channels that wrapped around the circular orientation space. ***E***, Decoding performance was quantified by subtracting the neural evidence at the presented orientation (e.g., 45°) from the evidence at the non-presented orientation (e.g., 135°). ***F***, As above, we collapsed across the presented gratings by subtracting the neural evidence for the presented (e.g., 45°) from the non-presented (e.g., 135°) orientation, for visualisation purposes. Shaded regions in B, C, E, and F indicate SEM.

**Figure 3.**
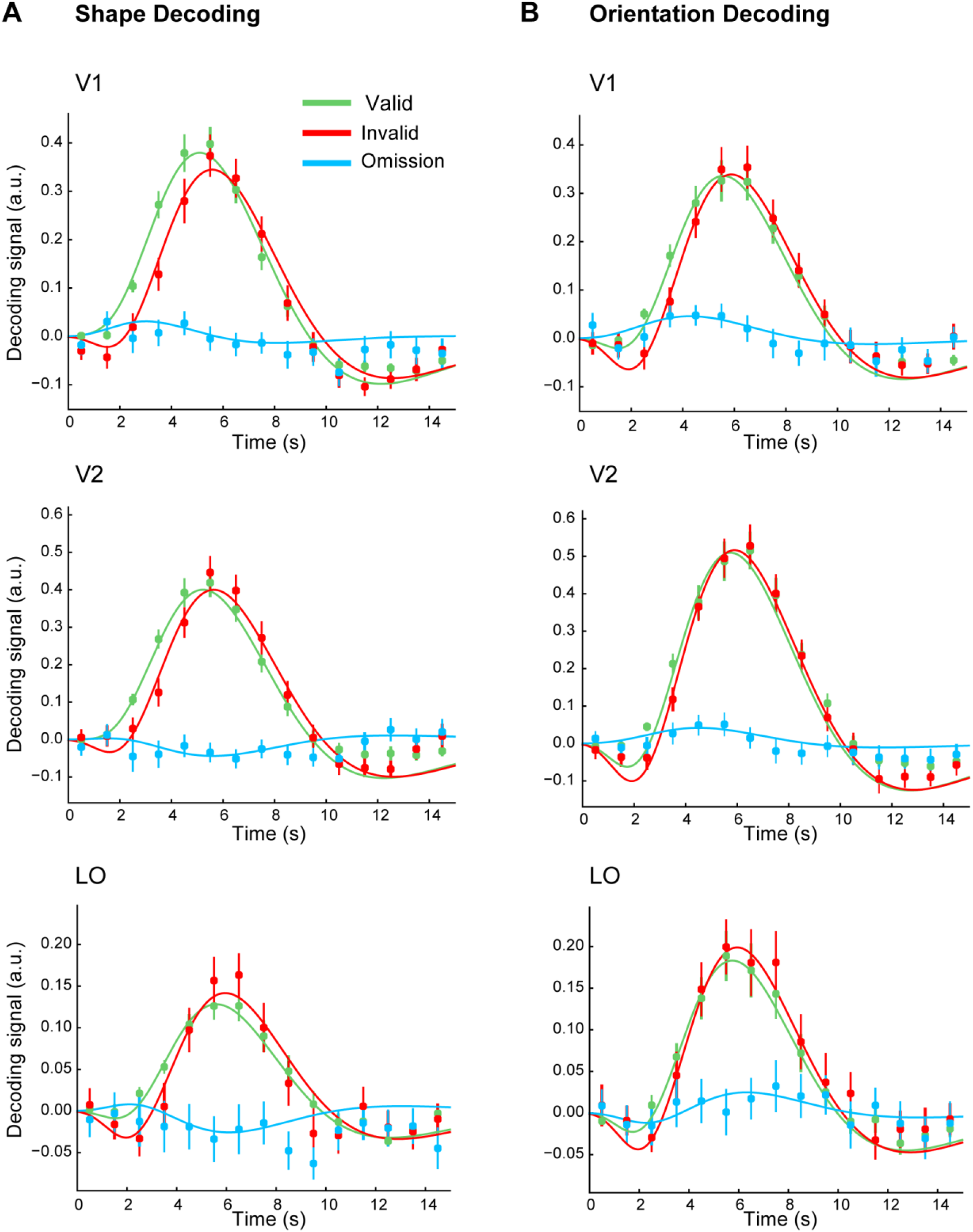
Time-resolved decoding in visual cortex for ***A***, complex shapes (Experiment 1) and B, oriented gratings (Experiment 2). Time course of the complex shape and grating orientation decoding signal separately for validly (green) and invalidly (red) predicted stimuli and predicted but omitted (blue) stimuli. The canonical HRF and its derivative were not fit to the fMRI data directly, but rather to the decoding signal obtained by reconstructing stimulus information for each time point from FIR parameter estimates. Error bars indicate SEM.

Valid predictions about complex shapes and grating orientations led to similar facilitation in visual cortex, across the cortical hierarchy (Fig. 3). Specifically, when the prediction was invalid, there was a delay in the decoding signal for shapes (V1: *t*_(23)_ = 3.40, *p* = 0.0024; V2: *t*_(23)_ = 3.06, *p* = 0.0056; marginal effect in LO: *t*_(23)_ = 1.96, *p* = 0.062) and gratings (V1: *t*_(23)_ = 2.87, *p* = 0.0086; V2: *t*_(23)_ = 2.15, *p* = 0.042; marginal effect in LO: *t*_(23)_ = 2.06, *p* = 0.051), relative to when it was valid. The peak of the decoding signal was also significantly lower for invalidly predicted shapes than for validly predicted shapes in V1 (*t*_23_ = 2.73, *p* = 0.012), but not V2 (*t*_23_ = 1.11, *p* = 0.27) or LO (*t*_23_ = 0.15, *p* = 0.87). There were no such effects for gratings across the visual hierarchy (V1: *t*_(23)_ = 1.27, *p* = 0.22; V2: *t*_(23)_ = 0.86, *p* = 0.40; LO: *t*_(23)_ < 0.01, *p* > 0.99). There were no significant differences in how predictions affected either the peak or the latency of the decoded stimulus signals between the two experiments (interaction between ‘prediction’ and ‘experiment’, all *p* > 0.3), consistent with the possibility that shape and orientation expectations modulate visual cortex processing similarly. In addition to the decoding signal, there was also a modest but highly reliable delay in the mean BOLD response evoked by invalidly compared to validly predicted shapes (V1: *t*_(23)_ = 6.33, *p* = 1.9 x 10 ^-6^; V2: *t*_(23)_ = 7.31, *p* = 1.9 x 10 ^-7^; LO: *t*_(23)_ = 7.48, *p* = 1.32 x 10 ^-7^) and gratings (V1: *t*_(23)_ = 5.73, *p* = 7.8 x 10^-6^; V2: *t*_(23)_ = 6.20, *p* = 3.2 x 10^-6^; LO: *t*_(23)_ = 5.38, *p* = 1.8 x 10^-5^). Prediction validity did not significantly affect the amplitude of the mean BOLD response in either experiment (all *p* > 0.05).

During the omission runs, the 25% non-valid trials did not involve the presentation of the unpredicted stimulus, but rather no visual stimulus at all. To investigate whether the BOLD response in visual cortex reflected the predicted (but omitted) stimuli, we inspected the fit of a canonical HRF to the decoding time course for these trials. This revealed that the orientation of expected but omitted gratings was successfully reconstructed from BOLD patterns in V1 (*t*_(23)_ = 2.28, *p* = 0.032), but not in V2 (*t*_(23_) = 1.37, *p* = 0.18) or LO (*t*_(23)_ = 0.59, *p* = 0.56), replicating Kok et al. (2014). However, expected but omitted shapes could not be reconstructed from BOLD patterns in any region (V1: *t*_(23)_ = 0.64, *p* = 0.53; V2: *t*_(23)_ = −1.33, *p* = 0.20; LO: *t*_(23)_ = −0.71, *p* = 0.49). Note that there were no reliable differences in decoding performance between omitted gratings and omitted shapes (V1: *t*_(46)_ = 1.06, *p* = 0.29; V2: *t*_(46)_ = 1.91, *p* = 0.062; LO: *t*_(46)_ = 0.92, *p* = 0.36).

### Hippocampus

Unlike visual cortex, complex shape and grating orientation predictions led to qualitatively different responses in the hippocampus (Fig. 4). In Experiment 1, the pattern of activity in the hippocampus contained a representation of the shape that was predicted by the auditory cue (*t*_(23)_ = 2.86, *p* = 0.0089; Fig. 4A), but was unaffected by the shape that was actually presented on screen (*t*_(23)_ = 0.54, *p* = 0.59). However, the situation was strikingly different for orientation in Experiment 2, where the pattern of activity in the hippocampus did not contain a representation of the orientation that was predicted by the cue (*t*_(23)_ = −1.59, *p* = 0.13; Fig. 4B). That is, there was a difference between shape and orientation prediction signals in the hippocampus (*t*_(46)_ = 3.04, *p* = 0.0039; Fig. 4C), driven by a positive shape prediction signal in Experiment 1 and a numerically negative orientation prediction signal in Experiment 2.

**Figure 4.**
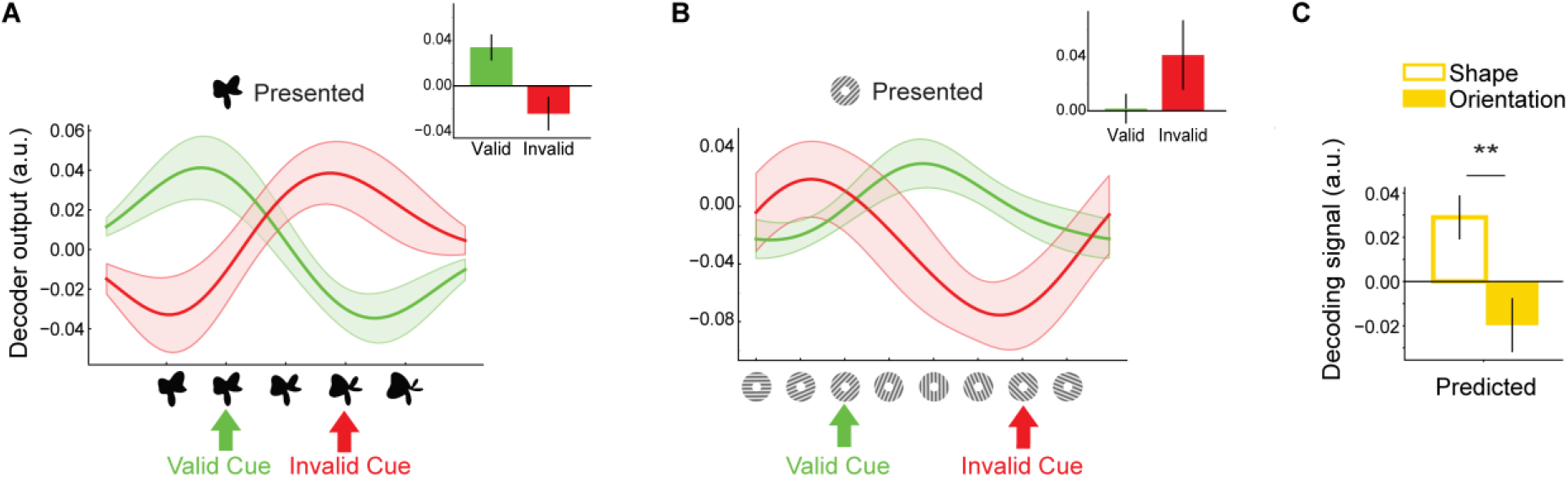
Stimulus reconstructions in hippocampus. ***A***, Shape reconstructions in the hippocampus were fully determined by the cued (predicted) shape, rather than the presented shape. ***B***, However, for grating orientations, when the cue was invalid, the hippocampus represented the unexpectedly presented orientation. ***C***, Predictions about complex shapes and grating orientation modulated the hippocampus differently. \*\**p* < 0.01. Shaded regions and error bars indicate SEM.

To interrogate the circuitry of these prediction signals further, we applied an automated segmentation method to define ROIs for the anatomical subfields of the hippocampus. Specifically, we segmented the hippocampus into CA1, CA2-3-DG, and subiculum. As in the hippocampus as a whole, representations of predicted shapes and orientations were strikingly different in CA2-3-DG (*t*_(46)_ = 3.07, *p* = 0.0036; Fig. 5C). Where Experiment 1 showed a trend towards evoking a *positive* representation of the expected shape in this subregion (*t*_(23)_ = 2.04, *p* = 0.053; Fig. 5A), grating expectations were *negatively* represented (*t*_(23)_ = −2.29, *p* = 0.031; Fig. 5B).

**Figure 5.**
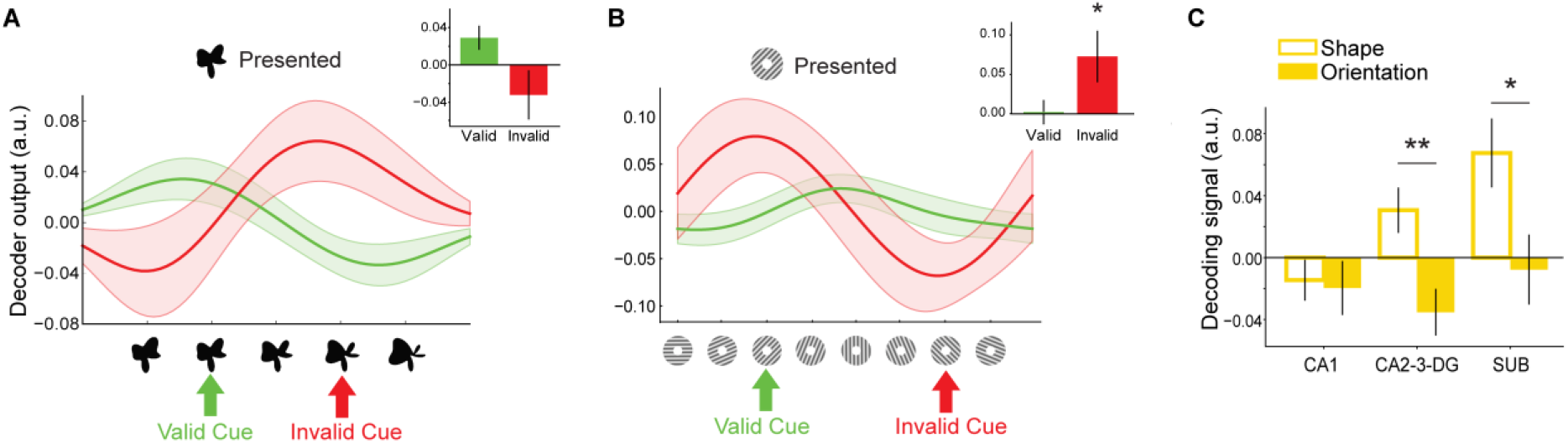
Stimulus reconstructions in CA2-3-DG. ***A***, As in hippocampus as a whole, shape reconstructions in the CA2-3-DG were fully determined by the cued (predicted) shape, rather than the presented shape. ***B***, However, for grating orientations, when the cue was invalid, CA2-3-DG represented the unexpectedly presented orientation. *C*, Decoding of the predicted stimuli across hippocampal subfields. \**p* < 0.05, \*\**p* < 0.01. Shaded regions and error bars indicate SEM.

To inspect these results more closely, we next compared the responses evoked by valid and invalid predictions in both experiments. In Experiment 1, when shape predictions were either valid and invalid, hippocampal representations were completely determined by the predicted shape. That is, when shape 2 was predicted and presented, hippocampal patterns represented shape 2, and when shape 4 was predicted but shape 2 was presented, hippocampus solely represented shape 4 (Fig. 4A, 5A). In Experiment 2, when grating orientation predictions were valid, activity patterns contained no evidence for any orientation, neither the predicted (and presented) orientation, nor the unpredicted orientation (hippocampus: *t*_(23)_ = 0.14, *p* = 0.89; CA2-3-DG: *t*_(23)_ = 0.13, *p* = 0.89; Fig. 4B, 5B). When orientation predictions were invalid, however, activity patterns reflected the (unexpectedly) presented orientation, most clearly in CA2-3-DG (*t*_(23)_ = 2.18, *p* = 0.04; hippocampus: *t*_(23)_ = 1.57, *p* = 0.13). In other words, only unexpectedly presented grating orientations were represented in CA2-3-DG, reminiscent of a prediction error type signal.

In the subiculum, predicted shapes and orientations evoked distinct representations as well (*t*_(46)_ = 2.32, *p* = 0.025; Fig. 5C). As in hippocampus as a whole, Experiment 1 revealed that shape representations in subiculum were dominated by the predicted shape (*t*_(23)_ = 2.97, *p* = 0.0069), but not the presented shape (*t*_(23)_ = −0.54, *p* = 0.59). In Experiment 2, on the other hand, neither predicted (*t*_(23)_ = −0.34, *p* = 0.74) nor presented (*t*_(23)_ = 0.52, *p* = 0.61) orientations were represented. That is, subiculum activity patterns did not contain information about grating orientation in any of the conditions. Finally, in CA1, neither predicted (*t*_(23)_ = −1.08, *p* = 0.29) or presented (*t*_(23)_ = 1.47, *p* = 0.15) shapes, nor predicted (*t*_(23)_ = −1.10, *p* = 0.28) or presented (*t*_(23)_ = 0.75, *p* = 0.46) orientations, were represented in the two experiments.

The subfield results were largely mimicked by representations evoked in omission trials. Expected but omitted shapes affected activity patterns in CA2-3-DG strikingly differently than expected but omitted orientations (*t*_(46)_ = 2.57, *p* = 0.013; Fig. 6). This difference was in the same direction as above, with shape prediction signals being more positive than orientation prediction signals. Note that the positive shape prediction signals (*t*_(23)_ = 1.64, *p* = 0.11) and negative orientation prediction signals (*t*_(23)_ = −2.00, *p* = 0.058) were not significant in isolation. Representations of expected but omitted stimuli did not significantly affect subiculum (shapes: *t*_(23)_ = 1.73, *p* = 0.097; orientations: *t*_(23)_ = −0.35, *p* = 0.73; difference: *t*_(46)_ = 1.47, *p* = 0.15) or CA1 (shapes: *t*_(23)_ = −0.46, *p* = 0.65; orientations: *t*_(23)_ = −0.67, *p* = 0.51; difference: *t*_(46)_ = 0.23, *p* = 0.82).

**Figure 6.**
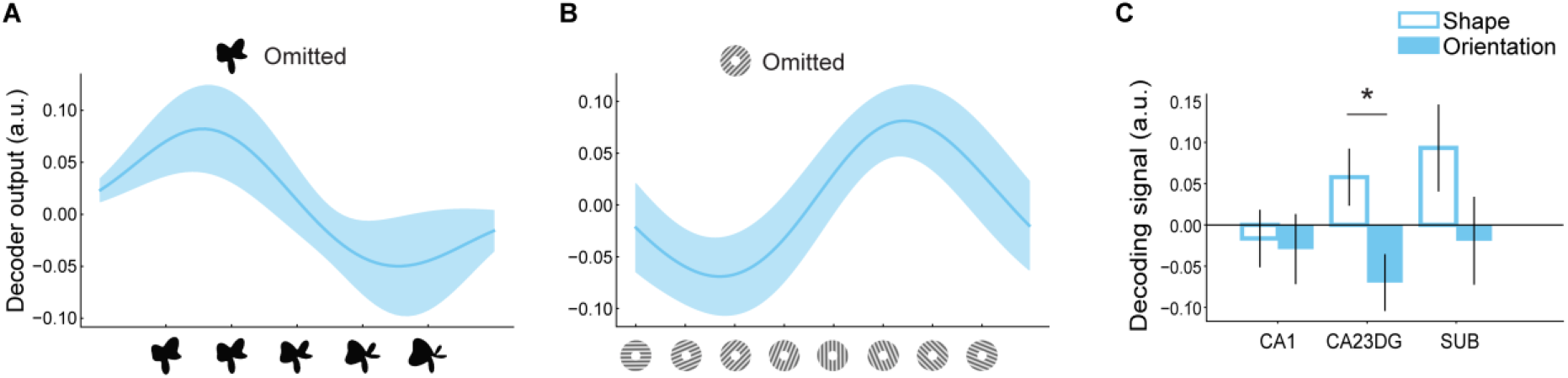
***A***, Reconstruction of expected but omitted shapes in CA2-3-DG. ***B***, Reconstruction of expected but omitted orientations in CA2-3-DG. C, Decoding of the expected but omitted stimuli across hippocampal subfields. \**p* < 0.05, \*\**p* < 0.01. Shaded regions and error bars indicate SEM.

Overall, the results for omission trials in the hippocampus resembled those for predicted stimuli when comparing valid and invalid trials. However, the effects were weaker statistically, perhaps due to lower signal-to-noise on trials without any visual stimulus or because of a qualitative difference between these conditions. An example of the latter could be that the omission of expected stimuli may trigger different cognitive processes than validly and invalidly cued trials. The absence of any visual stimulus is quite salient and surprising, given the regularity of their appearance in the rest of the study. In addition, participants did not perform a task on the omission trials, eliminating the need for perceptual decision making and response selection.

### Searchlight results

Our primary focus in this study was on the hippocampus, but to explore which other brain regions are involved in content-sensitive predictions, we performed searchlight analyses in the field of view of our functional scans (most of occipital and temporal and part of parietal and frontal cortex). This analysis revealed distinct representations of shape and orientation predictions in bilateral hippocampus, anterior occipital cortex, cerebellum, left inferior frontal gyrus, and left middle temporal gyrus, as well as a few smaller clusters elsewhere (Table 1). Separate searchlight analyses for the two experiments revealed positive representations of the predicted shape in the hippocampus, anterior occipital cortex, and a few smaller clusters, and negative orientation predictions in left inferior frontal gyrus, anterior occipital cortex and middle temporal gyrus (Table 1). The reverse contrasts (i.e., negative shape predictions and positive orientation predictions) did not reveal any significant clusters.

**Table 1.**
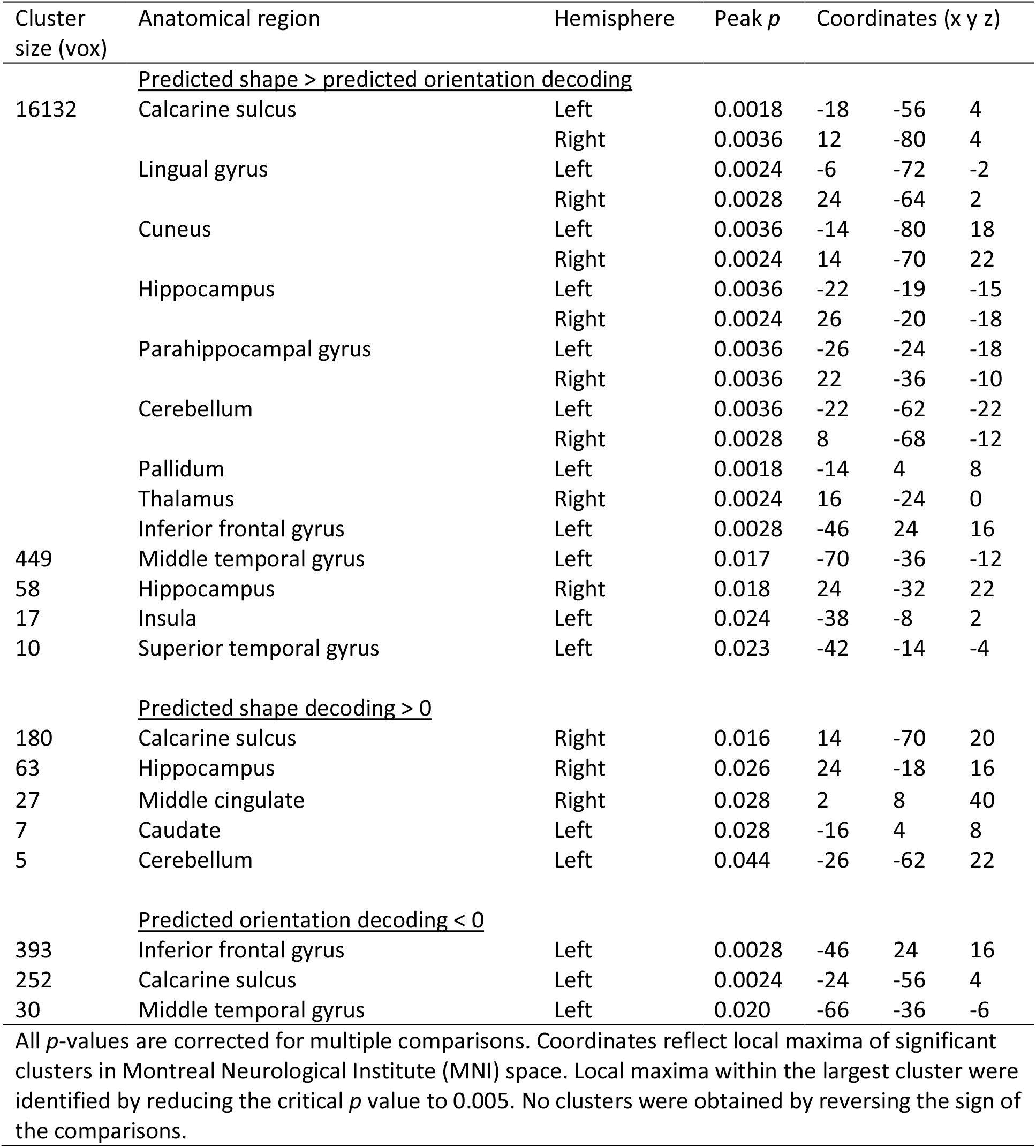
Searchlight results

### Control analyses

*Can the differences between shape and orientation experiments be explained by the fact that the two gratings associated with auditory cues were easier to distinguish than the two shapes associated with auditory cues (or vice versa)?* A difference in perceptual distance between the two shapes versus the two gratings may cause a different balance between pattern completion and pattern separation mechanisms in the hippocampus, thereby complicating our results. To examine this possibility, we split the participants in both experiments into two groups, depending on how well the two gratings or shapes could be decoded from V1, in the visual stimuli only runs. We then investigated whether hippocampal evidence for the predicted shape/orientation differed between high vs. low V1 decoders, using a two-way ANOVA. If the difference between the two experiments was driven by stimulus discriminability, we would expect to see a main effect of high vs. low decoders. However, we found a strong main effect of experiment (F_1,44_ = 9.79, *p* = 0.0031), no effect of high vs. low decoders (F_1,44_ = 1.24, *p* = 0.27), and no interaction between the two (F_1,44_ = 3.62, *p* = 0. 064). The trend towards an interaction reflected the fact that the positive prediction signal for shapes and the negative prediction signal for orientations tended to be stronger for the participants with *worse* V1 stimulus decoding. We are hesitant to interpret a marginal effect, but this could potentially reflect the need for stronger prediction in participants who had trouble disambiguating the stimuli. Regardless, this is in the opposite direction than hypothesized above, and thus does not provide evidence for an alternative explanation of content-sensitive hippocampal effects based on stimulus discriminability.

*Can the difference between the shape and orientation experiments be explained by a difference in behavioural task performance/difficulty?* Following the same logic as above, we split the participants in both experiments into two sub-groups based on their behavioural accuracy. We investigated whether evidence for the predicted shape/orientation in the hippocampus differed between high vs. low performers using a two-way ANOVA. This analysis revealed a main effect of experiment (*F*_1,44_ = 9.00, *p* = 0.0044), no main effect of task performance (*F*_1,44_ = 0.03, *p* = 0.87), and no interaction between the two (*F*_1,44_ = 0.87, *p* = 0.36). This fails to provide evidence that our main content-sensitive hippocampal effects can be attributed to differences in task difficulty.

*Can the difference between the shape and orientation experiments be explained by a difference in patterns of eye movements?* Another potential concern could be that the differential hippocampal effects could reflect predictive cues inducing different eye movements for shapes and gratings. In both experiments, participants were instructed to fixate on the bullseye in the center of the screen throughout the experiment and to not move their eyes towards the stimuli. Still, to examine potential influences of involuntary eye movements, we collected high quality eye-tracking data for 9 out of 24 participants in Experiment 1, and 15 out of 24 in Experiment 2. We investigated influences of the stimulus (i.e., shape 2 vs. shape 4 in Experiment 1; 45° vs. 135° grating in Experiment 2), predicted stimulus, and the interaction of the two, on pupil position both poststimulus (250-750 ms) and during the cue-stimulus interval (−250 to 0 ms).

We found no evidence of the presented shape on pupil position in Experiment 1, either prestimulus (x-coordinate: *t*_(8)_ = −0.14, *p* = 0.89; y-coordinate: *t*_(8)_ = 0.73, *p* = 0.49) or post-stimulus (x-coordinate: *t*_(8)_ = −1.44, *p* = 0.19; y-coordinate: *t*_(8)_ = −1.10, *p* = 0.30). In Experiment 2, there was a small difference in horizontal pupil position between 45° and 135° gratings post-stimulus (mean x-coordinate = 0.01° vs. −0.05°, respectively; *t*_(14)_ = 2.21, *p* = 0.044). This could reflect small involuntary eye-movements along the orientation axes of the gratings (40). There were no effects on prestimulus pupil position (x-coordinate: *t*_(14)_ = 1.95, *p* = 0.071; y-coordinate: *t*_(14)_ = −1.32, *p* = 0.21) or vertical post-stimulus pupil position (y-coordinate: *t*_(14)_ = −1.21, *p* = 0.25). However, crucial for the interpretation of our results is whether eye movements were influenced by the predictive cues. There were no effects of predicted shape, nor an interaction between predicted and presented shape, on pupil position in either pre- or post-stimulus intervals (all *p* > 0.10, both for horizontal and vertical pupil coordinates). Similarly, there were no effects of predicted orientation, nor an interaction between predicted and presented orientation, on pupil position in either pre- or poststimulus intervals (all *p* > 0.10, both for horizontal and vertical pupil coordinates). That is, we found no evidence for differences in eye movements that could explain the fMRI effects of the predictive cues. In addition to specifying pre- and post-stimulus time windows, we also conducted exploratory cluster-based permutation tests (41) on the full time window (−850 ms to 1000 ms). This analysis did not reveal any significant effects of presented or predicted stimulus, nor their interaction, for either Experiment 1 or 2 (no clusters *p* < 0.05).

## Discussion

Recent theories of the hippocampus suggest that it performs general-purpose computations independent of stimulus contents (20). Alternatively, it has been suggested that the nature of stimuli, especially their complexity, is a crucial factor in determining whether hippocampus and MTL are involved in a given perceptual task (31, 42). The current study addresses these hypotheses by revealing that predictions about complex shapes and simple grating orientations evoked qualitatively different representations in the hippocampus. This suggests that the hippocampus can play distinct computational roles in perception depending on the content of perceptual predictions, rather than executing a general-purpose process independent of stimulus content. This finding is especially noteworthy given that the experimental paradigms, fMRI scan sequences, and neural decoding methods were virtually identical in the two experiments. Furthermore, the effects of the predictions on processing in visual cortex were highly similar for complex shapes and oriented gratings, suggesting that the hippocampal differences were not due to simple differences in the extent to which the predictions were learned or used to guide perception.

Could our results have been caused by something other than the nature of the stimuli per se? For instance, it could be that the two cue-associated shape stimuli were harder to distinguish from one another than the two cue-associated gratings (or vice versa), changing the computations in hippocampus, e.g., affecting the balance between pattern completion and separation. However, in a control analysis, we did not find any evidence of stimulus discriminability driving the difference in hippocampal effects between the two experiments. Another possibility is that differences in task difficulty between the two experiments (detecting a subtle shape morph vs. detecting a subtle grating phase shift) drove the hippocampal differences. However, an adaptive staircasing procedure ensured that task performance was roughly matched between experiments and a control analysis did not reveal evidence of task performance driving hippocampal effects. Finally, despite clear instructions to keep their eyes fixated, it is conceivable that participants moved their eyes in response to the predictive cues and did so differently for the two experiments, leading to differences in hippocampal responses. Aside from the fact that this might be expected to also lead to differences in visual cortex between the two experiments, analysis of the eye-tracking data did not reveal any effects of the predictive cues on gaze position, in either experiment.

In sum, the differential responses of the hippocampus in the two experiments seem best explained by the difference in the nature of the predicted stimuli: complex objects versus simple features. This is in line with theories on hierarchical message passing in the brain (26–28). Complex objects are known to be represented in the MTL, such as in perirhinal cortex (31, 32), areas known to have direct reciprocal connections with the hippocampus (17, 21). Therefore, the hippocampus is ideally positioned to supply complex shape predictions to its immediate cortical neighbours. Specifically, upon reception of a predictive cue (rising or falling tones), pattern completion mechanisms in hippocampus, especially in the CA3 subfield (11, 16, 43), may lead to retrieval of the predicted associate, which can then be sent back to MTL cortex as a prediction of upcoming inputs. In contrast, for the low-level feature of orientation, hippocampus does not seem to represent the predicted feature. This may be explained by the fact that the nearby cortical recipients of hippocampal feedback in the MTL do not preferentially represent such low-level features. Rather, as reviewed above, these areas represent complex objects abstracted away from their simple features (i.e., invariant over location, size, etc.) and are thus not ideal targets for predictions about such features.

In fact, for oriented gratings, the hippocampus and especially its CA3 subfield (combined with CA2 and dentate gyrus) seemed to represent prediction *errors* rather than predictions (44–46): validly cued orientations were cancelled out whereas invalidly cued orientations were not. In other words, the hippocampus seemed to represent an ‘anti-prediction’ that inhibited representation of expected stimuli. This is consistent with the observed negative evidence for expected but omitted orientations. Such coding for stimulus prediction errors may allow the hippocampus to refine learning and predictions elsewhere in the brain. Note that we observed these effects in CA2-3-DG, whereas most theories propose that prediction errors or at least a comparison between retrieved and experienced information should occur in CA1 (44, 45). It is possible that we did not observe such CA1 effects because we scanned after associative learning of the cues and outcomes was complete, and that they may be more apparent if we had examined responses during the learning process, a possibility that awaits future studies.

Interestingly, shape prediction signals were strongly present in the subiculum, but orientation signals were fully absent there. The subiculum is a major output hub of hippocampus back to MTL cortex (17, 47). Therefore, this pattern of results is in line with the suggestion that hippocampus may be a top-down source for high-level object predictions in visual cortex, but not for low-level feature predictions. This proposal leads to distinct hypotheses about the direction of signal flow through the hippocampal and MTL system during processing of complex shape and feature predictions, respectively. That is, shape predictions are proposed to flow from the hippocampus (CA3 through subiculum) back to cortex via entorhinal cortex (EC), whereas orientation predictions are not (and prediction errors may flow forward from EC to CA1/CA3). These hypotheses can be tested in future research using layer-specific fMRI of EC (48, 49), since signals flowing into hippocampus arise from superficial layers, whereas signals flowing from hippocampus back to cortex arrive in the deep layers (17). Additionally, this can be addressed using simultaneous electrophysiological measurements in hippocampus and cortex, for instance in human epilepsy patients, which offer superior temporal resolution.

When oriented gratings were expected but omitted, the pattern of activity in V1 reflected the expected orientation, suggesting that such expectations can evoke a template of the predicted feature in sensory cortex, in line with previous findings (3, 39). In contrast, this did not occur for expected but omitted shapes. Together with the differential hippocampal representations, these findings suggests that expectations about low-level features and higher-level objects may involve distinct neural mechanisms.

The effects of the orientation predictions on grating-evoked signals in visual cortex, as reported here, differ from those reported previously using a similar paradigm (2). Whereas invalid grating orientation predictions in that study led to both an increased peak BOLD amplitude and a reduced orientation representation in V1 (2), the current study found that invalid orientation predictions lead to *delayed* signals, both in terms of BOLD amplitude and orientation representations. Although the cause of this difference is currently unclear, there were a couple of potentially important differences between these studies. First, the two studies employed different behavioural tasks: participants performed orientation and contrast discrimination in Kok et al. (2012a), whereas in the current study they discriminated grating phase. Although we do not have a clear hypothesis for how this would lead to differences in V1, previous work has shown that task demands influence expectation effects in visual cortex (50–53). Additionally, there were differences in the grating presentations between the two studies. In Kok et al. (2012a), the individual gratings were presented for a longer duration (500 vs. 250 ms) but with a shorter inter-stimulus interval (100 vs. 500 ms), and the two gratings in a given trial were in anti-phase with different spatial frequencies. Whether and how these parameters could explain the differential effects of expectation cues on V1 processing is unclear, but one possibility is that they might affect the degree of repetition suppression between the two gratings in each trial, which could in turn interact with prediction signals (2, 54–56).

In sum, the current study revealed that predictions about complex shapes and simple orientations evoke distinct representations in hippocampus. These findings are in line with the hippocampus generating perceptual predictions for high-level objects, but not for low-level features. This fits well with hierarchical Bayesian inference theories of sensory processing (26–28), which suggest that each brain region provides predictions to those regions with which it has direct feedback connections, and formats those predictions in the currency that the receiving region ‘understands’ (27, 57). Finally, these findings suggest that stimulus complexity is a crucial factor in determining whether, and in what role, the hippocampus is involved in perceptual inference (31).

## Materials and Methods

### Participants

Experiment 1 enrolled 25 healthy individuals with normal or corrected-to-normal vision. Participants provided informed consent to a protocol approved by the Princeton University Institutional Review Board and were compensated ($20 per hour). One participant was excluded from analysis because they moved their head between runs so much that their occipital lobe was partly shifted outside the field of view. The final sample consisted of 24 participants (15 female, mean age 23). We previously reported some findings from this dataset (12), though we performed additional analyses for present purposes that are reported here.

Experiment 2 enrolled 24 healthy individuals with normal or corrected-to-normal vision (15 female, mean age 24). Participants provided informed consent to a protocol approved by the Yale University Human Investigation Committee and were compensated ($20 per hour). This is a new dataset not previously reported.

### Stimuli

Visual stimuli were generated using MATLAB (Mathworks; RRID:SCR_001622) and the Psychophysics Toolbox (58; RRID:SCR_002881). In both experiments, stimuli were displayed on a rear-projection screen using a projector (1920 x 1068 resolution, 60 Hz refresh rate) against a uniform grey background. Participants viewed the stimuli through a mirror mounted on the head coil. Auditory cues consisted of three pure tones (440, 554, and 659 Hz; 80 ms per tone; 5ms intervals), presented in ascending or descending pitch through headphones.

In Experiment 1, the visual stimuli were complex shapes defined by radio frequency components (RFCs) (59, 60, 61; Fig. 1A-D). These stimuli were created by varying seven RFCs and were based on a subset of the stimuli used by Op de Beeck et al. (2001; see their Fig. 1a). By varying the amplitude of three of the seven RFCs, a one-dimensional shape space was created. Specifically, the amplitudes of the 1.11, 1.54, and 4.94 Hz components increased together, ranging from 0 to 36 (first two components) and from 15.58 to 33.58 (third component). Five shapes were chosen along this continuum to create a perceptually symmetrical set, centred on the third shape (for details, see 12). To create slightly warped versions of the shapes, in order to enable a same/different discrimination task, a fourth RFC (the 3.18 Hz component) was modulated. The shape stimuli were presented in black (subtending 4.5°), centred on a fixation bullseye.

In Experiment 2, visual stimuli consisted of grayscale luminance-defined sinusoidal gratings that were displayed in an annulus (outer diameter: 10°, inner diameter: 1°, spatial frequency: 1.5 cycles/°), surrounding a fixation bullseye (Fig. 1E-H). Eight gratings were used to span the 180° orientation space, in equal steps of 22.5°. To enable a similar same/different discrimination task as in Experiment 1, we modulated the phase of the gratings.

### Experimental procedure

Each trial of Experiment 1 started with the presentation of a fixation bullseye (0.7°). During the ‘prediction’ runs, an auditory cue (ascending or descending tones, 250 ms) was presented 100 ms after onset of the trial. After a 500-ms delay, two consecutive shape stimuli were presented for 250 ms each, separated by a 500-ms blank screen (Fig. 1A). The auditory cue (ascending vs. descending tones) predicted whether the first shape on that trial would be shape 2 or shape 4, respectively (out of five shapes; Fig. 1B). The cue was valid on 75% of trials, while in the other 25% of trials the unpredicted shape would be presented. For instance, an ascending auditory cue might be followed by shape 2 on 75% of trials and by shape 4 on the remaining 25% of trials. During omission runs, the cues were also 75% valid, but on the remaining 25% of trials no shape was presented at all, with only the fixation bullseye remaining on screen. All participants performed two prediction runs (128 trials, ~13 min per run) and two omission runs (128 trials, ~13 min per run), in interleaved ABBA fashion (order counterbalanced across participants). Halfway through the experiment, the contingencies between the auditory cues and the shapes were flipped (e.g., ascending tones were now followed by shape 4 and descending by shape 2). The order of the cue-shape mappings was counterbalanced across participants. Participants were trained on the cue-shape associations during two practice runs (112 trials total, ~8 min) in the scanner, one before the first prediction/omission run and one halfway through the experiment after the contingency reversal. During these practice runs, the auditory cue was 100% predictive of the identity of the first shape on that trial (e.g., ascending tones were always followed by shape 2 and descending tones by shape 4). The two practice runs took place while anatomical scans (see below) were acquired, to make full use of scanner time. On each trial, the second shape was either identical to the first or slightly warped. This warp was achieved by modulating the amplitude of the orthogonal 3.18 Hz RFC component defining the shape by an amount much smaller that the differences between shape indices on the continuum defined over the three other varying components. This modulation could be either positive or negative (counterbalanced across conditions) and participants’ task was to indicate whether the two shapes on a given trial were the same or different.

The design of Experiment 2 was identical to Experiment 1, except that the visual stimuli consisted of oriented gratings instead of complex shapes (Fig. 1E-H). That is, Experiment 2 contained two prediction and two omission runs, but here the auditory cues (rising and falling tones, 250 ms) predicted the orientation of an upcoming grating (45° or 135°, 250 ms). A second grating (250 ms) with the same orientation was presented after a 500-ms delay, and was either identical or slightly phase shifted with respect to the first grating. As with the shape modulation for the same/different task, changes in phase were orthogonal and much smaller than the differences in orientation across stimuli. Participants’ task was to indicate whether the two gratings were the same or different.

Finally, both experiments contained two additional runs in which no auditory cues were presented. Each trial started with the presentation of the fixation bullseye, a delay of 850 ms (to equate onset with runs containing the auditory cues), the first stimulus (250 ms), another delay of 500 ms, and the second stimulus (250 ms). The two visual stimuli (Experiment 1: complex shapes, Experiment 2: oriented gratings) were either identical or slightly different (Experiment 1: warped shape, Experiment 2: phase shift). Participants indicated whether the two visual stimuli were the same or different. In Experiment 1 (120 trials, ~13 min per run), each trial contained one of the five shapes with equal (20%) likelihood (Fig. 1D). In Experiment 2 (128 trials, ~13 min per run), each trial contained one of the eight gratings with equal (12.5%) likelihood (Fig. 1H). These ‘non-predictive’ runs were designed to be as similar as possible to the prediction/omission runs, save the absence of the predictive auditory cues. They were collected as the first and last runs of each session and the data were used to train the neural decoding models (see below).

In both experiments, participants indicated their response using an MR-compatible button box. After the response interval ended (750 ms after disappearance of the second visual stimulus), the fixation bullseye was replaced by a single dot, signalling the end of the trial while still requiring participants to fixate. Also in both experiments, the magnitude of the difference between the two stimuli on a given trial (Experiment 1: shape warp, Experiment 2: phase offset) was determined by an adaptive staircasing procedure (62), updated after each trial, in order to make the same/different task challenging (~75% correct) and comparable in difficulty across experiments. Separate staircases were run for trials containing valid and invalid cues, as well as for the non-predictive runs, to equate task difficulty between conditions. The staircases were kept running throughout the experiments. They were initialised at a value determined during an initial practice session 1-3 days before the fMRI experiment (no auditory cues, 120 trials). After the initial practice run, the meaning of the auditory cues was explained, and participants practiced briefly with both cue contingencies (valid trials only; 16 trials per contingency). Because participants were practicing both mappings equally, this session did not serve to train them on any particular cue-shape association. Rather, this session was intended to familiarize participants with the structure of trials and the nature of the experiment.

### MRI Acquisition

For Experiment 1, structural and functional data were collected using a 3T Siemens Prisma scanner with a 64-channel head coil at the Princeton Neuroscience Institute. Functional images were acquired using a multiband echoplanar imaging (EPI) sequence (TR, 1000 ms; TE, 32.6 ms; 60 transversal slices; voxel size, 1.5 × 1.5 × 1.5mm; 55° flip angle; multiband factor, 6). This sequence produced a partial volume for each participant, parallel to the hippocampus and covering most of the temporal and occipital lobes. Anatomical images were acquired using a T1-weighted MPRAGE sequence (TR, 2300 ms; TE, 2.27 ms; voxel size, 1 × 1 × 1 mm; 192 sagittal slices; 8° flip angle; GRAPPA acceleration factor, 3). Two T2-weighted turbo spin-echo (TSE) images (TR, 11,390 ms; TE, 90 ms; voxel size, 0.44 × 0.44 × 1.5 mm; 54 coronal slices; perpendicular to the long axis of the hippocampus; distance factor, 20%; 150° flip angle) were acquired for hippocampal segmentation. To correct for susceptibility distortions in the EPI, a pair of spin-echo volumes was acquired in opposing phase-encode directions (anterior/posterior and posterior/anterior) with matching slice prescription, voxel size, field of view, bandwidth, and echo spacing (TR, 8000 ms; TE, 66 ms).

For Experiment 2, data were acquired on a 3T Siemens Prisma scanner with a 64-channel head coil at the Yale Magnetic Resonance Research Centre. Functional images were acquired using a multiband EPI sequence with virtually identical parameters to Experiment 1 (TR, 1000 ms; TE, 33.0 ms; 60 transversal slices; voxel size, 1.5 × 1.5 × 1.5mm; 55° flip angle; multiband factor, 6), as was the pair of opposite phase-encode spin-echo volumes for distortion correction (TR, 8000 ms; TE, 66 ms). Anatomical images were similar to Experiment 1. T1-weighted images were acquired using an MPRAGE sequence (TR, 1800 ms; TE, 2.26 ms; voxel size, 1 × 1 × 1 mm; 208 sagittal slices; 8° flip angle; GRAPPA acceleration factor, 2). Two T2-weighted TSE images were acquired (TR, 11,170 ms; TE, 93ms; voxel size, 0.44 × 0.44 × 1.5 mm; 54 coronal slices; distance factor, 20%; 150° flip angle).

### fMRI preprocessing

Images for both experiments were preprocessed using FEAT 6 (FMRI Expert Analysis Tool), part of FSL 5 (http://fsl.fmrib.ox.ac.uk/fsl, Oxford Centre for Functional MRI of the Brain, RRID:SCR_002823) (63). All analyses were performed in participants’ native space. Using FSL’s topup tool (64), susceptibility-induced distortions were determined on the basis of opposing-phase spin-echo volumes. This output was converted to radians per second and supplied to FEAT for B0 unwarping. The first six volumes of each run were discarded to allow T1 equilibration, and the remaining functional images for each run were spatially realigned to correct for head motion. These functional images were registered to each participant’s T1 image using boundary-based registration and temporally high-pass filtered with a 128 s period cut-off. No spatial smoothing was applied. Lastly, the two T2 images were co-registered and averaged, and the resulting image was registered to the T1 image through FLIRT (FMRIB’s Linear Image Registration Tool).

### Regions of interest

Our main focus was the hippocampus. Using the automatic segmentation of hippocampal subfields (ASHS) machine learning toolbox (65) and a database of manual medial temporal lobe segmentations from a separate set of 51 participants (66, 67), hippocampal regions of interest (ROIs) were defined based on each participant’s T2 and T1 images for CA2-CA3-DG, CA1, and subiculum subfields. CA2, CA3, and DG were combined into a single ROI because these subfields are difficult to distinguish with fMRI. Results of the automated segmentation were visually inspected for each participant to ensure accuracy.

In visual cortex, ROIs were defined for V1, V2, and lateral occipital (LO) cortex in each participant’s T1 image using Freesurfer (http://surfer.nmr.mgh.harvard.edu/; RRID:SCR_001847). To ensure that we were measuring responses in the retinotopic locations corresponding to our visual stimuli, we restricted the visual cortex ROIs to the 500 most active voxels during the non-predictive runs. Since no clear retinotopic organization is present in the hippocampal ROIs, cross-validated feature selection was used instead (see below). All ROIs were collapsed over the left and right hemispheres, since we had no hypotheses regarding hemispheric differences.

### fMRI data modeling

Functional data were modeled with general linear models (GLM) using FILM (FMRIB’s Improved Linear Model). This included temporal autocorrelation correction and extended motion parameters (six standard parameters, plus their derivatives and their squares) as nuisance covariates.

For Experiment 1, we specified regressors for the conditions-of-interest: shape-only runs, five shapes; prediction runs, two shapes x two prediction conditions (valid vs invalid); omission runs, two shapes x two omission conditions (presented vs omitted). Delta functions were inserted at the onset of the first shape (or expected onset, for omissions) of each trial and convolved with a double-gamma hemodynamic response function (HRF). The same procedure was used for Experiment 2, with the following convolved regressors: grating only runs, eight orientations; prediction runs, two orientations x two prediction conditions (valid vs invalid); omission runs, two orientations x two omission conditions (presented vs omitted). For both experiments, we also included the temporal derivative of each regressor to accommodate variability in the onset of the response (37).

Additional finite impulse response (FIR) models were fit in both experiments to investigate the temporal evolution of shape and grating representations in visual cortex. This approach estimated the BOLD signal evoked by each condition-of-interest at 20 x 1 s intervals. We trained the decoder on the FIR parameter estimates from the shape-only or grating-only runs, averaging over the time points spanning 4-7 s (corresponding to the peak hemodynamic response). This decoder was then applied to the FIR parameter estimates from all of the time points in the prediction and omission runs. The amplitude and latency of this time-resolved shape/grating information was quantified by fitting a double gamma function and its temporal derivative to the decoder output.

### Decoding analysis

In both experiments, we used a forward modelling approach to reconstruct visual stimuli from the pattern of BOLD activity in a given brain region (36). This approach has proven successful in reconstructing continuous stimulus features, such as hue (36), orientation (68), and motion direction (69).

In our case, the continuous dimensions of shape contour and grating orientation were modelled by a set of hypothetical channels, each with an idealised tuning curve (i.e., basis function). The difference between the analyses in Experiments 1 and 2 was entirely in the definition of these hypothetical channels. To decode shape in Experiment 1, the basis functions consisted of five hypothetical channels, each a halfwave-rectified sinusoid raised to the fifth power, spaced evenly such that they were centred on the five points in shape space that constituted the five shapes presented in the experiment (Fig. 2A). To decode orientation in Experiment 2, the basis functions consisted of six halfwave-rectified sinusoids raised to the fifth power spaced evenly along the 180° orientation space (Fig. 2D). The number of orientation basis functions was based on previous studies decoding circular stimulus features (36, 68, 69). Note that the orientation space, unlike the shape space, was circular, and therefore the channels wrapped around from 180° to 0°.

Other than the hypothetical channels/basis functions, the forward modelling approach was identical in Experiments 1 and 2. In the first ‘training’ stage, BOLD activity patterns obtained from the shape-only or grating-only runs were used to estimate the weights of these channels for each voxel, using linear regression. Specifically, let *k* be the number of channels, *m* the number of voxels, and *n* the number of measurements (i.e., the five shapes or the eight orientations in the non-predictive runs). The matrix of estimated BOLD response amplitudes for the different stimuli (**B**_train_, *m x n)* was related to the matrix of hypothetical channel outputs (C_train_, *k x n)* by a weight matrix (**W**, *m x k*):

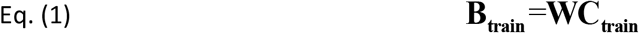

The least-squares estimate of this weight matrix W was estimated using linear regression:

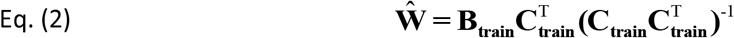

These weights reflected the relative contribution of the hypothetical channels in the forward model to the observed response amplitude of each voxel. Using these weights, the second stage of analysis reconstructed the channel outputs associated with the test patterns of activity across voxels evoked by the stimuli in the main experiment (i.e., the prediction and omission runs; **B**_test_), again using linear regression. This step transformed each vector of *n* voxel responses (parameter estimates per condition) into a vector of *k* channel responses. These channel responses (C_test_) were estimated using the learned weights (**W**):

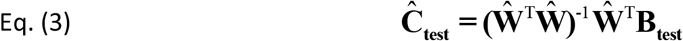

The channel outputs were used to compute a weighted average of the basis functions, reflecting neural evidence over the shape or orientation dimension (Fig. 2B,E). During the prediction and omission runs of Experiment 1, only shapes 2 and 4 were presented (Fig. 1B). Thus, four neural evidence curves were obtained for these runs: two shapes by two prediction/omission conditions (valid vs. invalid/presented vs. omitted). We collapsed across the presented shape by subtracting the neural evidence for shape 4 from that for shape 2, thereby subtracting out any non-shape-specific BOLD signals (Fig. 2C). Analogously, in Experiment 2, we subtracted the neural evidence for 135° gratings from that for 45° gratings (Fig. 2F).

For statistical testing, scalar decoding performance values were calculated on the basis of decoded neural evidence. For Experiment 1, decoding performance during the prediction/omission runs was quantified by subtracting the neural evidence for the presented shape (e.g., shape 2) from that of the non-presented shape (e.g., shape 4). For Experiment 2, decoding performance was quantified by subtracting the neural evidence for the presented orientation (e.g., 45°) from that of the non-presented orientation (e.g., 135°). (Note: For the omission runs there was no presented shape or orientation, and so we conditioned decoding performance on the *expected* shape or orientation.) For each participant in the two experiments, this procedure led to a measure of decoding performance for validly and invalidly predicted stimuli (shapes or gratings), respectively. This allowed us to quantify evidence for the stimuli as presented on the screen (by averaging evidence for validly and invalidly predicted stimuli) and evidence for the cued stimuli [by averaging (1 – evidence) for the invalidly predicted stimuli with evidence for the validly predicted stimuli]. Finally, we calculated decoding performance for predicted but omitted stimuli (shapes and gratings, respectively). These measures were statistically tested at the group level using *t* tests.

In visual cortex ROIs, we selected the 500 most active voxels during the shape- and grating-only runs. However, the hippocampus does not show a clear evoked response to visual stimuli. Therefore, we applied a different method of voxel selection for hippocampal ROIs. Voxels were first sorted by their informativeness, that is, how different the weights for the forward model channels were from each other. Second, the number of voxels to include was determined by selecting between 10 and 100% of the voxels, increasing in 10% increments. We then trained and tested the model on these voxels within the shape- and grating-only runs (trained on one run and tested on the other). For all iterations, decoding performance on shapes 2 and 4 (Experiment 1) or 45° and 135° orientations (Experiment 2) was quantified as described above, and we selected the number of voxels that yielded the highest performance (group average: Experiment 1, 1536 of 3383 voxels; Experiment 2, 1369 of 3229 voxels). We also labelled the selected hippocampus voxels based on their subfield from segmentation (group average: Experiment 1, 436 voxels in CA1; 572 voxels in CA2-CA3-DG; 425 voxels in subiculum; Experiment 2, 394 voxels in CA1; 490 voxels in CA2-CA3-DG; 380 voxels in subiculum).

For the main ROI and searchlight analyses, the input to the forward model consisted of parameter estimates from a voxelwise GLM that fit the amplitude of the BOLD response using regressors convolved with a double-gamma HRF. However, we also supplied parameter estimates from a data-driven FIR model that made no assumptions about the timing or shape of BOLD responses.

### Searchlight

A multivariate searchlight approach was used to explore the specificity of predicted shape and orientation representations, as well as significant differences between the two, within the field of view of our functional scans (most of occipital and temporal and part of parietal and frontal cortex). In both experiments, a spherical searchlight with a radius of 5 voxels (7.5mm) was passed over all functional voxels, using the searchlight function implemented in the Brain Imaging Analysis Kit (BrainIAK, http://brainiak.org, RRID:SCR_014824). In each searchlight for each participant, we performed shape/orientation decoding in the same manner as in the ROIs, which yielded maps of decoding evidence for the predicted shapes (Experiment 1) and orientations (Experiment 2).

Each participant’s output volumes were registered to the MNI 152 standard template for group analysis. This was achieved by applying the non-linear registration parameters obtained from registering each participant’s T1 image to the MNI template using AFNI’s (RRID:SCR_005927) 3dQwarp (https://afni.nimh.nih.gov/pub/dist/doc/program_help/3dQwarp.html). Group-level nonparametric permutation tests were applied to these searchlight maps using FSL randomise (70), correcting for multiple comparisons using threshold-free cluster enhancement (71). To determine where in the brain orientation and shape prediction signals differed from one another, we conducted a 2-sample t-test comparing predicted shape evidence to predicted orientation evidence at *p* < 0.05 (two-sided). This test yielded one large cluster, so we identified local maxima within the cluster by reducing the critical *p* value to 0.005. Follow-up one-sample t-tests used to explore the two experiments separately at *p* < 0.05 (one-sided).

### Experimental design and statistical analysis

The two experiments were designed to compare differences in BOLD signals evoked by predictions of complex shapes and grating orientations, respectively. For hippocampal ROIs, we quantified decoding evidence for presented, predicted, and omitted stimuli (shapes and orientations). For visual cortex ROIs, we compared differences in both the amplitude and the latency of the decoding signal as a result of valid vs. invalid predictions. For details on how these values were computed, see the ‘Decoding analysis’ section above. Our primary interest was the differential effects of prediction for shape vs. orientation. Such effects were tested using independent-samples t-tests, as shape and orientation predictions were measured in different participant groups. Within the two experiments, effects of predictions were tested using paired-sample t-tests (e.g., valid vs. invalid). Statistical testing of the whole-brain searchlight is described above in the ‘Searchlight analysis’ section.

### Data and code accessibility

Data and code are available upon request from the first author.

## Acknowledgements

This work was supported by an NWO Rubicon grant 446-15-004 to P.K. and NIH R01 MH069456 to N.B.T.-B. The Wellcome Centre for Human Neuroimaging is supported by core funding from the Wellcome Trust (203147/Z/16/Z).

## Author contributions

P.K. and N.B.T.-B. designed the experiment, P.K. and L.I.R. conducted the experiment, P.K. and L.I.R. analysed the data, P.K., L.I.R. and N.B.T.-B. wrote the paper.

